# Investigating the potential of X shredding for mouse genetic biocontrol

**DOI:** 10.1101/2023.12.05.570030

**Authors:** Mark. D. Bunting, Gelshan I. Godahewa, Nicole O. McPherson, Louise Robertson, Luke Gierus, Sandra G. Piltz, Owain Edwards, Mark Tizard, Paul Q. Thomas

## Abstract

CRISPR-Cas9 technology has facilitated development of strategies that can potentially provide more humane and effective methods to control invasive vertebrate species, such as mice. One promising strategy is X chromosome shredding which aims to bias offspring towards males, resulting in a gradual and unsustainable decline of females. This method has been explored in insects with encouraging results. Here, we investigated this strategy in *Mus musculus* by targeting repeat DNA sequences on the X chromosome with the aim of inducing sufficient DNA damage to specifically eliminate X chromosome-bearing sperm during gametogenesis. We tested three different guide RNAs (gRNAs) targeting different repeats on the X chromosome, together with three male germline-specific promoters for inducing Cas9 expression at different stages of spermatogenesis. A modest bias towards mature Y-bearing sperm was detected in some transgenic males, although this did not translate into significant male-biasing of offspring. Instead, cleavage of the X-chromosome during meiosis typically resulted in a spermatogenic block, manifest as small testes volume, empty tubules, low sperm concentration, and sub/infertility. Our study highlights the importance of controlling the timing of CRISPR-Cas9 activity during mammalian spermatogenesis and the sensitivity of spermatocytes to X chromosome disruption.

## Introduction

Invasive species pose a significant risk to biodiversity and are responsible for a global economic cost of $423 billion, including the associated reduction in quality of life^1^ for people affected. Current pest management practices are often expensive, ineffective, have ethical considerations, and can cause death of non-target species, either from direct consumption of baits or ingestion of bait-containing prey^2^. Consequently, there is a pressing and unmet need for innovative strategies in invasive species control that offer enhanced efficacy and specificity.

With the advent of CRISPR-Cas9 genome editing technology, genetic biocontrol strategies to modify or supress invasive species and disease vector populations have become an area of intense interest. Synthetic gene drive systems to spread an inactivated fertility gene through a population have shown promise in Dipteran insects, but remain challenging to adapt to rodents^3–5^. The recent development of *t*-CRISPR, an engineered version of the naturally-occurring *t* haplotype male meiotic drive that carries a CRISPR cassette targeting a haplosufficient female fertility gene, provides the first and only validated strategy for mammalian genetic biocontrol^6^. An alternative suppression strategy involves biasing the sex of offspring through disruption of either Y- or X-bearing sperm, increasing the relative proportion of females or males, respectively. This sex-distorter technique, termed X-shredding, has been successful in insects^7–10^. Experimentation from our lab, and others, has shown *in silico, in vitro*, and *in vivo* feasibility of deleting the Y chromosome^11–13^. The use of gRNAs and Cas9 to generate XO zygotes through deletion of an X chromosome in females has also been demonstrated^13^. Chromosomal elimination is achieved using gRNAs that target repeat sequences found only within the target chromosome. When combined with Cas9, multiple double-stranded breaks are induced along repeat region which can either be resolved through non-homologous end joining (NHEJ) repair (generating a local deletion), loss of the target chromosome arm, or the entire chromosome. Spatial modelling of X chromosome shredding drives in rodents has shown their potential for the eradication of mouse populations on islands^14^.

Here, we build on the concept of X shredding using transgenic mice. Using a safe split drive format, we tested a range of male germline-specific promoters driving Cas9 expression in combination with gRNAs targeting different X chromosome repeats. Despite robust DNA cutting activity and induction of deletions, overt male bias was not observed. This study identifies obstacles that must be overcome to successfully develop X-shredder technology in rodents.

## Results

### In vitro gRNA-mediated cutting of the X chromosome

We initially tested X-shredder gRNAs in mouse embryonic stem (mES) cells to determine the efficiency of X chromosome elimination. Mouse ES cells transfected with gRNA constructs (X-B, X-C, X-D^13^) showed significant cell death compared to control gRNAs that target a single copy autosomal gene (Tyrosinase) or without a genomic target (Neomycin), supporting X chromosome shredding activity (normalised to Tyr 100%, X-B 63.8 ± 5.3%, X-C 57.1 ± 14.7%, X-D 3.4 ± 0.12%, X-B+X-C 27.5 ± 24.8%, X-B+X-D 1.1 ± 0.3%, X-C+X-D 1.1 ± 0.59%) (Fig. 1A,B). We found that X-D resulted in the greatest level of cell death (>95%) compared with X-B (>35%) and X-C (>40%). Dual gRNA expression (X-B+X-C, X-B+X-D, and X-C+X-D) further increased cell lethality. Sequencing of target regions revealed indels at single cut sites in remaining cells, confirming DNA cleavage at target sites (Suppl. Fig. 1). We further validated DNA cutting activity of gRNAs by scoring DNA double strand breaks (DSB) detected via gamma-H2AX antibody binding (Fig. 1C,D). Mouse ES cells transfected with X-shredder gRNA showed significantly higher DSBs (3.3-4.6 fold) (five-fold) compared with the tyrosinase gRNA control, further supporting X-shredder gRNA cleavage activity. The mCherry gRNA does not target mouse genome, consistent with the near absence of DSBs.

**Figure 1.**
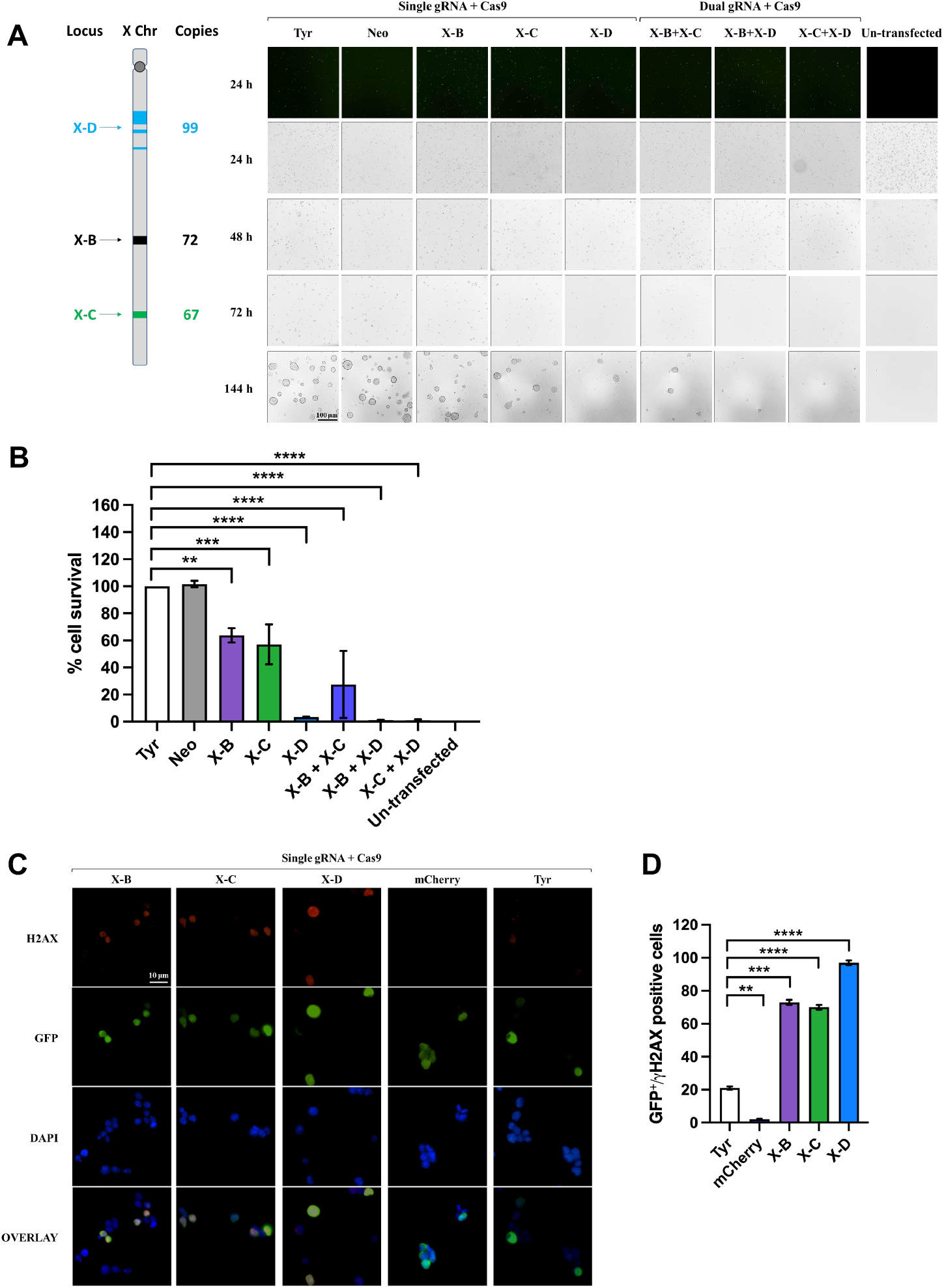
Elimination of the X chromosome and creation of DNA double strand break in mouse ES cells by CRISPR/Cas9-mediated gene editing. **A**. Schematic of X-B, X-C, and X-D target sites and their copies on the mouse X chromosome and representative microscopy images of mouse ES cells 24 h, 48 h, 72 h, and 144 h after transfection. Scale bar = 100 μM. **B**. Cell survival percentage at 144 h post-transfection of X-shredder gRNAs (n=3; Bars show mean ± SD, one way ANOVA with Sidak’s multiple comparison test). **C**. Representative immunofluorescence images of mouse ES cells 24 h after transfection. Scale bar = 10 μM. **D**. Average numbers of GFP/gamma-H2AX double positive cells (n=3). Bars show mean ± SD, one way ANOVA with Sidak’s multiple comparison test.

### X-shredder transgenic mouse generation

To assess X-shredding activity *in vivo*, we employed a ‘split drive’ system. gRNA- and Cas9-expressing mouse lines were generated separately and then crossed, resulting in ubiquitous gRNA and male germline-specific Cas9 expression. All transgenes were randomly integrated into the genome using pronuclear injection. We established Cas9 expression lines using the previously validated germline promoter sequences of *Ccna1, Prm1*, and *Stra8*^*15–19*^ which are active in pre-meiotic, meiotic (leptotene-zygotene) and post-meiotic spermatocytes, respectively. Cas9 was linked to an EGFP reporter using a P2A self-cleaving peptide (Suppl. Fig. 2A;^6^). X-B, X-C, and X-D gRNA sequences were driven by the U6 promoter. A CMV mCherry fluorescent reporter cassette was also included to facilitate identification of transgene carriers (Suppl. Fig. 2B). We generated single lines for X-B and X-D which contained 40 and 10 transgene copies, respectively (Fig. 2A). Two independent X-C lines, X-C-1 and X-C-2 were generated, both of which carried eight copies of the transgene (Fig. 2A). All lines were positive for mCherry fluorescence as determined by ear skin biopsies, although, X-D mice had much lower expression compared with the other lines (Fig. 2B).

**Figure 2.**
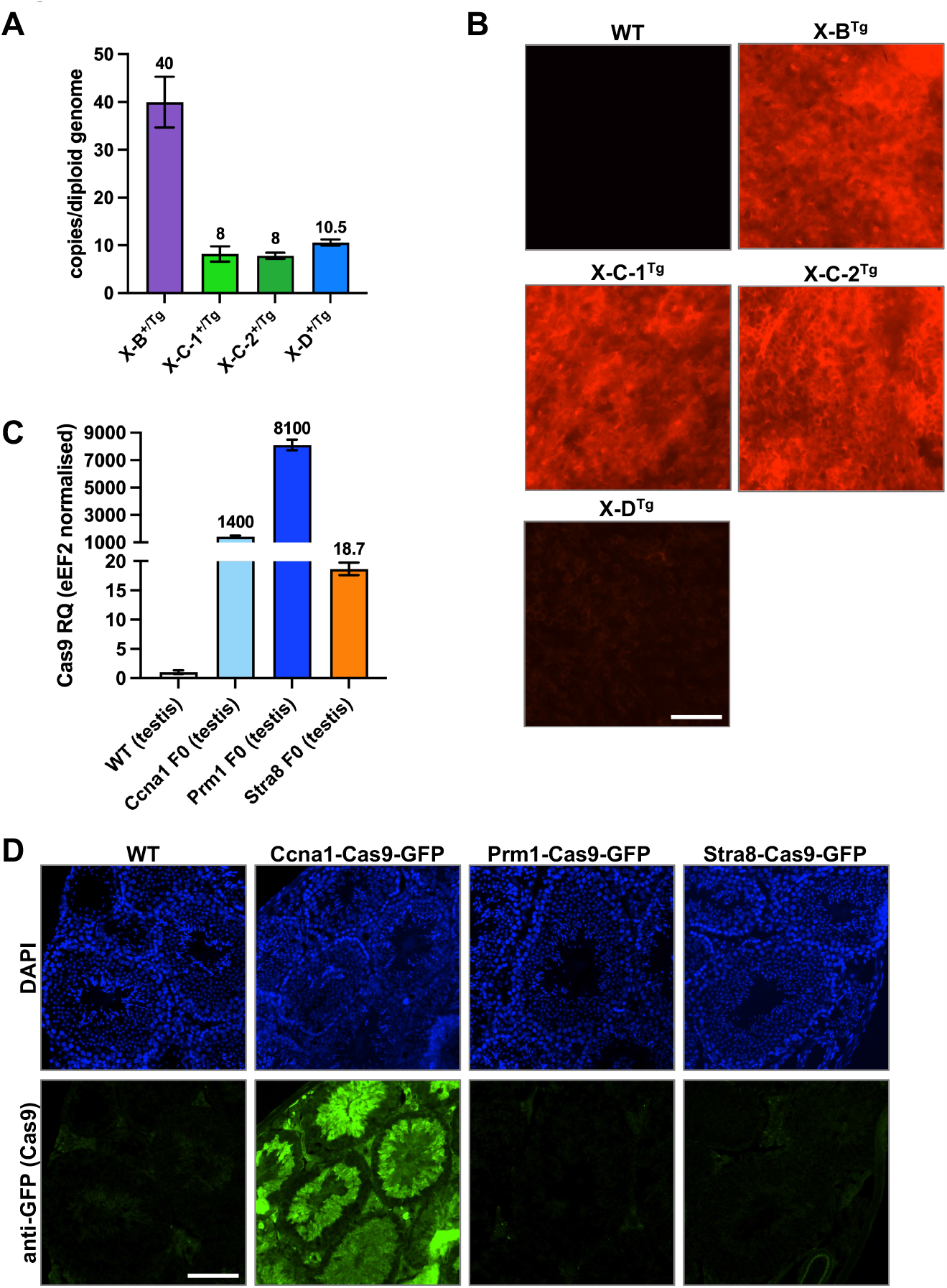
Characterisation of X shredder-mCherry and germline promoter-Cas9 transgenic mice. **A**. Transgene copy number of hemizygous X-B (n = 2), X-C-1 (n = 3), X-C-2 (n = 3) (two founder lines), and X-D (n = 10) mice. Bars show mean ± SD. **B**. Fluorescence imaging performed on ear skin punch biopsies for the transgenic mouse lines described in A. showing representative mCherry signal (red) with a WT mCherry negative control. *n* = >30 per genotype. Scale bar = 200 μM. **C**. Expression of Cas9 RNA in testis isolated from *Ccna1*-, *Prm1*-, and *Stra8*-Cas9 transgenic mouse lines. Expression is normalised to *eEF2* and WT testis indicates background Cas9 detection in this assay. *n* = 1 per genotype. **D**. Representative IF of Cas9-GFP expression (green) in the testis of *Ccna1*-Cas9-GFP transgenic mice. GFP signal was amplified by staining with an anti-GFP antibody while DAPI nuclear staining (blue) shows tubule structures. n = 2 per genotype. Scale bar = 100 μM.

RNA harvested from the testis of Cas9-EGFP transgenic mouse lines was used for qRT-PCR to determine *Cas9* expression levels. The *Ccna1*-Cas9-EGFP and *Prm1*-Cas9-EGFP lines expressed high levels of *Cas9* mRNA while lower transgene expression was detected in *Stra8*-Cas9-EGFP testes (Fig. 2C). However, we could only detect EGFP fluorescence in testis sections of *Ccna1*-Cas9-EGFP mice, where it localised to the tubules and was expressed in late spermatocytes (D and m) to elongating spermatids (1-14 of spermiogenic cycle) (Fig. 2D, Suppl. Fig. 2C). We therefore focussed mainly on the Ccna1-Cas9 line for *in vivo* experiments.

### Characterisation of gRNA; Cas9 double transgenic mice

We generated double transgenic progeny expressing an X-shredding gRNA together with Cas9 by mating X-B, X-C (two lines, X-C-1 and X-C-2), or X-D lines with *Ccna1*-Cas9-EGFP. The weight of *Ccna1*^Tg^; X-B^Tg^ testis were comparable to wild type (WT), while both *Ccna1*^Tg^; X-C^Tg^ lines and *Ccna1*^Tg^; X-D^Tg^ mice had significantly smaller testis (Fig. 3A). Histological analysis showed relatively normal tubule structure in *Ccna1*^Tg^; X-B^Tg^ mice while both *Ccna1*^Tg^; X-C^Tg^ lines and *Ccna1*^Tg^; X-D^Tg^ mice exhibited disrupted tubule architecture (Fig. 3B). Two-dimensional area and perimeter of *Ccna1*^Tg^; X-C^Tg^ and *Ccna1*^Tg^; X-D^Tg^ testes were also significantly decreased compared with WT (Fig. 3C and D, respectively). Tubules either lacking or with few spermatogenic cells were common in *Ccna1*^Tg^; X-C^Tg^ and *Ccna1*^Tg^; X-D^Tg^ testes (Fig. 3B,E,F), as were the presence of tubule vacuoles. All double transgenic lines showed significantly reduced sperm concentrations and dramatically reduced sperm motility fitness compared with WT (Fig. 3G,H).

**Figure 3.**
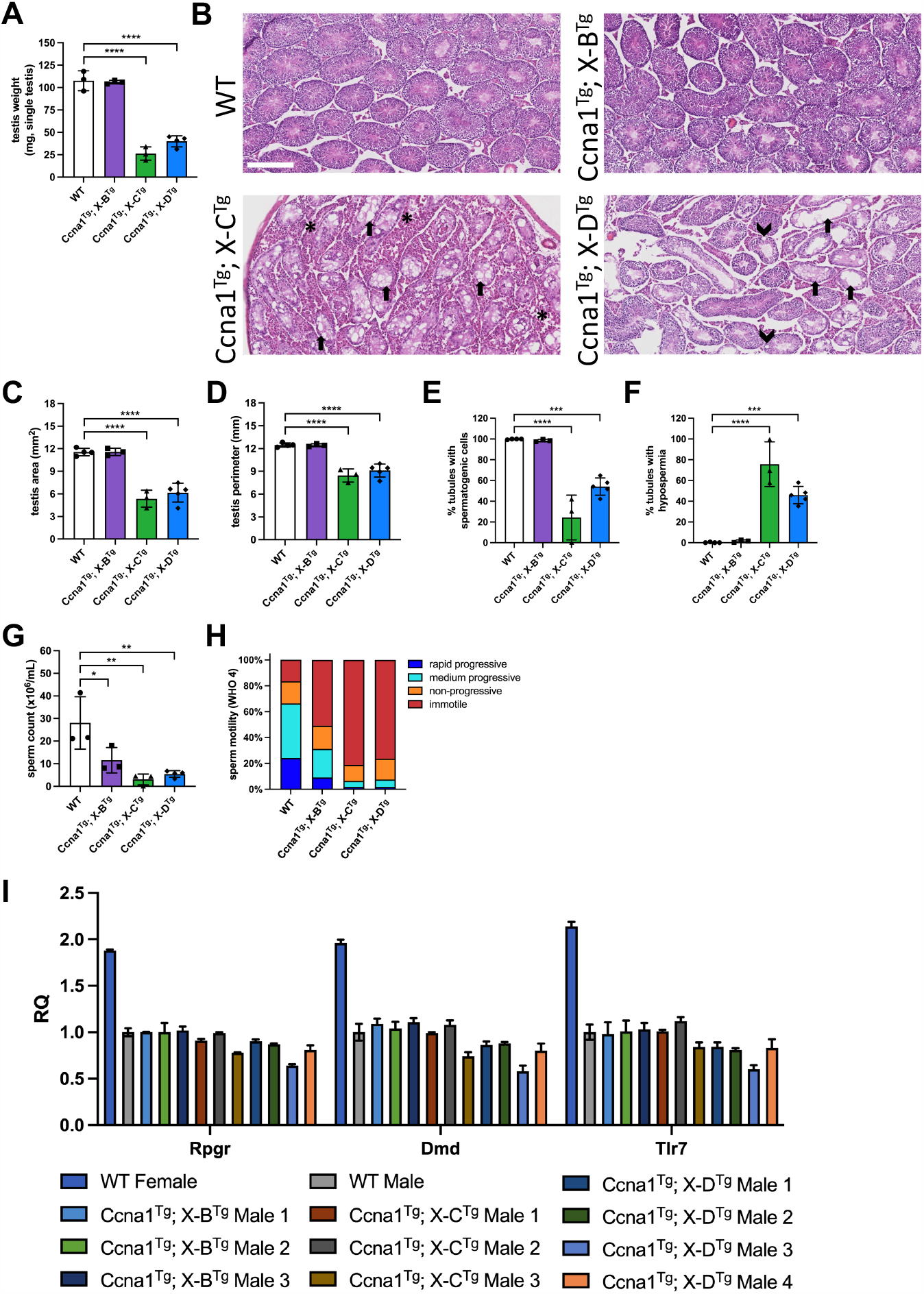
Functional assessment of *Ccna1*-Cas9^Tg^; X shredder^Tg^ double transgenic mice. **A**. WT, *Ccna1*^Tg^; X-B^Tg^, *Ccna1*^Tg^; X-C^Tg^, and *Ccna1*^Tg^; X-D^Tg^ ex-breeder males were sacrificed, testis removed and weighed on a fine balance. **B**. H&E staining of representative testis sections from mice used in A. showing disrupted tubule architecture in *Ccna1*^Tg^; X-C^Tg^ and *Ccna1*^Tg^; X-D^Tg^ mice. Arrows indicate empty tubules. Arrow heads indicate vacuoles. Asterisks indicate areas with an abundance of Leydig cells. Scale bar = 250 μM. Quantification of two-dimensional area **C**. and perimeter size **D**. from sections. **E**. Enumeration of the frequency of tubules containing spermatogenic cells. **F**. Quantification of the frequency of empty tubules. **G**. Sperm isolated from the epididymis were enumerated using a Sperm Class Analyser. **H**. Sperm motility was measured according to WHO 4 metrics. Mean motility values from mice shown. **I**. X chromosome dosage was assessed by qPCR using sperm DNA isolated from WT, *Ccna1*^Tg^; X-B^Tg^, *Ccna1*^Tg^; X-C^Tg^, and *Ccna1*^Tg^; X-D^Tg^ mice. Female mouse DNA containing two X chromosomes was used as the reference. **A**. and **C-H**. n = 3 for WT, *Ccna1*^Tg^; X-B^Tg^, *Ccna1*^Tg^; X-C^Tg^, and n = 4 *Ccna1*^Tg^; X-D^Tg^. Bars show mean ± SD. **A, C-I** show X-C-1^Tg^ and X-C-2^Tg^ data combined. Bars show mean ± SD, one way ANOVA with Sidak’s multiple comparison test.

To investigate if disproportionate X chromosome sperm loss was occurring *in vivo*, we isolated sperm from the vas deferens and cauda epididymis of WT and double transgenic male mice and measured X chromosome dosage by qPCR using three genes spanning the X chromosome (*Rpgr, DMD*, and *Tlr7*). Interestingly, *Ccna1*^Tg^; X-D^Tg^ male 3 and *Ccna1*^Tg^; X-C^Tg^ male 3 had lower gene dosage at all three loci (Fig. 3L). The former was particularly striking with only ∼60% X chromosome dosage detected. *Ccna1*^Tg^; X-D^Tg^ males 1, 2 and 4 also had lower dosage across 2 of the 3 loci. In contrast, *Ccna1*^Tg^; X-B^Tg^ males had normal X chromosome dosage. These data suggest that disproportionate X chromosome loss is occurring *in vivo* when the X-C and X-D gRNA and Cas9 are expressed in the male germline, although the effect is variable and incompletely penetrant.

### Biased generation of male offspring is not observed in experimental matings

To investigate male offspring bias, we mated *Ccna1*-Cas9^Tg^; X-shredder-gRNA^Tg^ males with WT females and assessed the proportion of male and female progeny (Fig. 4A). Cas9^Tg^; gRNA^Tg^ females (in which the spermatogenesis promoters driving Cas9 expression would not be active) were used as controls. Progeny from *Ccna1*^Tg^; X-B^Tg^ and *Ccna1*^Tg^; X-C^Tg^ sires from both X-C^Tg^ lines did not show significant deviation from a 50:50 male:female ratio (Fig. 4B,C,D). Similarly, Cas9 expression driven by *Prm1* or *Stra8* did not significantly alter the male:female ratio when used in combination with the both X-C^Tg^ lines (Fig. 4E,F). Interestingly, *Ccna1*^Tg^; X-D^Tg^ males sired an excess of male progeny (152 males versus 122 females) but this failed to reach statistical significance (*P* = 0.0699). To further assess male bias, we performed *in vitro* fertilisation using sperm isolated from a *Ccna1*^Tg^; X-D^Tg^ male. Following generation of zygotes, blastocysts were allowed to develop, DNA was extracted, and the ratio of male to female blastocysts was enumerated. No significant difference in the male:female ratio was observed, confirming that significant loss of X-bearing sperm was not generally occurring in double transgenic males (Suppl. Fig. 3B).

**Figure 4.**
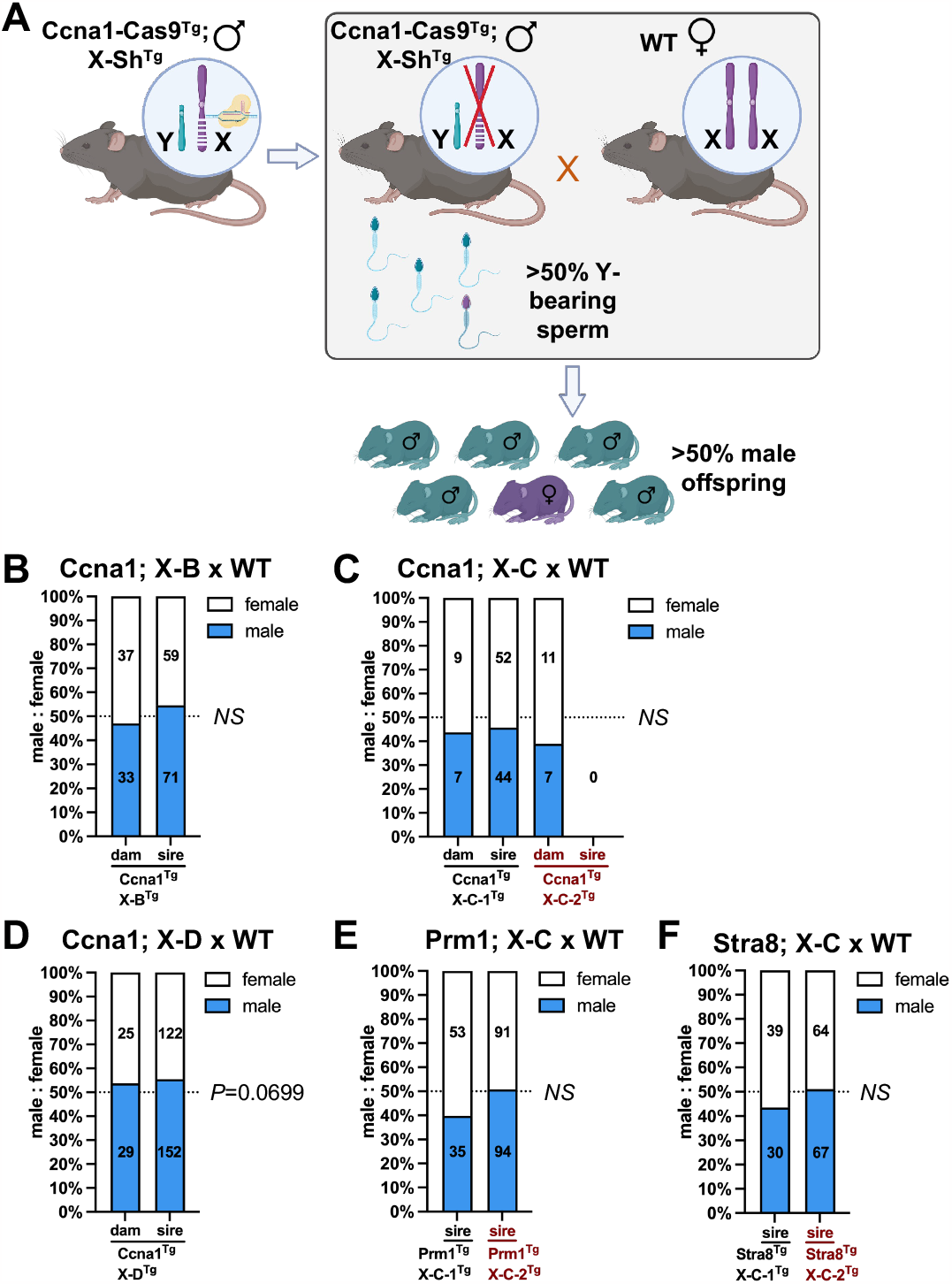
Transgenic mice co-expressing X shredder gRNAs and germline promoter-Cas9 did not generate altered male bias. **A**. Schematic showing the method to determine if male bias was occurring in offspring of germline promoter-Cas9^Tg^; X shredder^Tg^ male x WT females. Control crosses consisted of germline promoter-Cas9^Tg^; X shredder^Tg^ females x WT males. Enumeration of female and male pups from: **B**. *Ccna1*^Tg^; X-B^Tg^ x WT matings. Control matings = 11 plugs from three transgenic females with 10 litters, experimental matings = 29 plugs from three transgenic males with 22 litters. **C**. *Ccna1*^Tg^; X-C^Tg^ x WT matings (two independent X-C^Tg^ founder lines). *Ccna1*^Tg^; X-C-1^Tg^ control matings = 4 plugs from one transgenic female with 3 litters, *Ccna1*^Tg^; X-C-1^Tg^ experimental matings = 28 plugs from one transgenic male with 15 litters. *Ccna1*^Tg^; X-C-2^Tg^ control matings = 3 plugs from one transgenic female with 3 litters, *Ccna1*^Tg^; X-C-2^Tg^ experimental matings = 29 plugs from two transgenic males with 0 litters. **D**. *Ccna1*^Tg^; X-D^Tg^ x WT matings. Control matings = 7 plugs from three transgenic females with 8 litters, experimental matings = 109 plugs from five transgenic males with 48 litters. **E**. *Prm1*^Tg^; X-C^Tg^ x WT matings (two independent X-C^Tg^ founder lines). *Prm1*^Tg^; X-C-1^Tg^ experimental matings = 16 plugs from one transgenic male with 14 litters. *Prm1*^Tg^; X-C-2^Tg^ experimental matings = 42 plugs from three transgenic males with 27 litters. **F**. *Stra8*^Tg^; X-C^Tg^ x WT matings (two independent X-C^Tg^ founder lines). *Stra8*^Tg^; X-C-1^Tg^ experimental matings = 16 plugs from one transgenic male with 10 litters. *Stra8*^Tg^; X-C-2^Tg^ experimental matings = 20 plugs from three transgenic males with 18 litters. **B-F**. Numbers in bars indicate number of female pups (white) and male pups (blue) combined from litters of the same genotype. Chi squared test.

### Fertility defects in gRNA; Cas9 double transgenic mice

Given the small testes and low sperm count in some double transgenic males, we compared litter size and plug-to-pregnancy rates with controls (double transgenic females, where available) and historical non-transgenic pregnancy rates from WT mice maintained in our animal facility. No significant changes in litter size were seen in double transgenic mice compared with single transgenic controls kept under the same conditions (Suppl. Fig. 2A-E). *Ccna1*^Tg^; X-B^Tg^, *Prm1*^Tg^; X-C-1^Tg^, *Prm1*^Tg^; X-C-2^Tg^, *Stra8*^Tg^; X-C-1^Tg^ and *Stra8*^Tg^; X-C-2^Tg^ double transgenic males sired litters at normal rates (Suppl. Fig. 2A,C,D). The *Ccna1*^Tg^; X-C-1^Tg^ line exhibited normal fertility in contrast to the *Ccna1*^Tg^; X-C-2^Tg^ line which failed to have any litters (Suppl. Fig. 2B). *Ccna1*^Tg^; X-D^Tg^ double transgenic males varied in their ability to sire pups, with 2/5 males exhibiting normal fertility, while 3/5 had reduced fertility (Suppl. Fig. 2E).

### Triple transgenic X shredder mice are infertile

Given that the gRNA target sites for X-B, X-C and X-D are relatively localised on the X chromosome, we reasoned that expressing two different gRNAs targeting independent repeats would increase the likelihood of generating irreparable damage, resulting in X chromosome loss. Therefore, we generated *Ccna1*^Tg^; X-B^Tg^; X-C^Tg^ (Fig. 5A, X-C-1 and X-C-2 lines), *Ccna1*^Tg^; X-B^Tg^; X-D^Tg^ (Fig. 5B), and *Ccna1*^Tg^; X-C^Tg^; X-D^Tg^ (Fig. 5C, X-C-1 and X-C-2 lines) triple transgenic males which were mated with WT females to assess male offspring bias. However, triple transgenic males were unable to sire litters with WT dams despite successful plug generation. Female triple transgenic mice had expected pregnancy success (Fig. 5A-C, left) and generated litters with normal male:female ratios (Fig. 5A-C, right). Histological examination of the testis from triple transgenic males revealed severe disruption to tubule architecture and lack of any spermatogenic cells, consistent with azoospermia and their inability to sire offspring (Fig. 5D).

**Figure 5.**
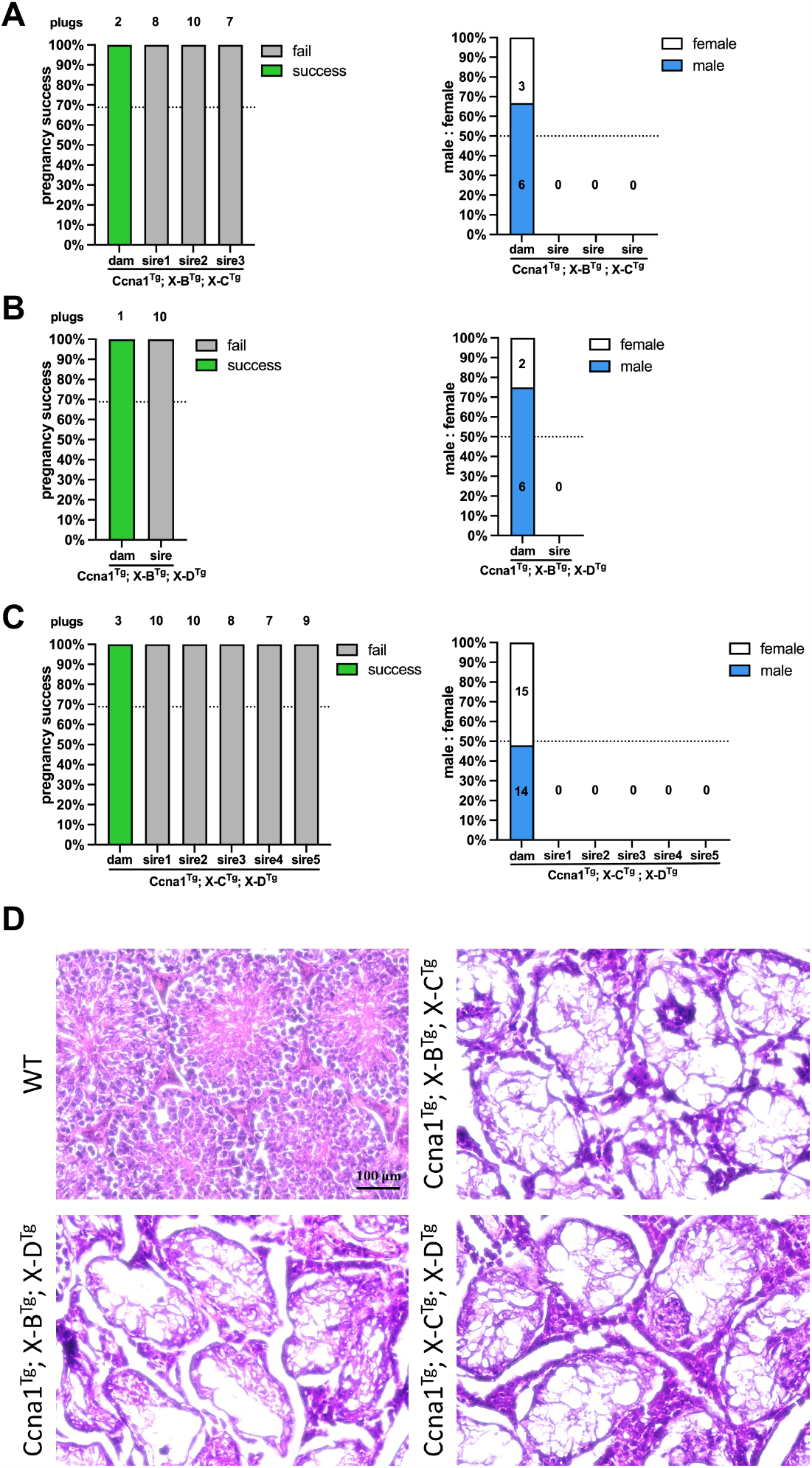
Functional assessment of Ccna1-Cas9^Tg^; X shredder^Tg^ triple transgenic mice. A-C pregnancy success as determined by plug and successful pregnancy counts (left) and enumeration of male and female pups (right) from: **A**. *Ccna1*^Tg^; X-B^Tg^; X-C^Tg^ x WT matings. Control matings = 2 plugs from one transgenic female with 2 litters, experimental matings = 25 plugs from three transgenic males with zero litters. **B**. *Ccna1*^Tg^; X-B^Tg^; X-D^Tg^ x WT matings. Control matings = 1 plug from one transgenic female with 1 litter, experimental matings = 10 plugs from one transgenic male with zero litters. **C**. *Ccna1*^Tg^; X-C^Tg^; X-D^Tg^ x WT matings. Control matings = 3 plugs from one transgenic female with 3 litters, experimental matings = 44 plugs from five transgenic males with zero litters. *D*. H&E staining of representative testis sections from mice used in A-C. Scale bar = 100 μM.

### Molecular characterisation of in vivo X chromosome cleavage

To better understand the cleavage activity of X-shredder gRNAs during spermatogenesis, we looked for indels at gRNA target sites in female offspring from double transgenic sires. We focussed initially on *Ccna1*^Tg^; X-B^Tg^ offspring, as the target repeats are contained within a localised 1,800 bp region that is amenable to PCR amplification (Fig. 6A). Analysis of the X-B target site from 37 randomly selected females showed large deletions of varying sizes (Fig. 6A). Sequencing of the PCR products and alignment to the WT sequence confirmed the presence of large deletions in all the sequenced samples, indicating robust cleavage of the X-B target site (Fig. 6A). As the repeat sequences targeted by X-C and X-D span much larger regions, a single target site was selected for a similar analysis of 13 randomly selected females from *Ccna1*^Tg^; X-C-1^Tg^ sires and 31 from *Ccna1*^Tg^; X-D^Tg^ sires. One sample from *Ccna1*^Tg^; X-C-1^Tg^ sires and 2 samples from *Ccna1*^Tg^; X-D^Tg^ sires had indels at the selected single cut site showing a low rate of indels (Fig. 6B). No target site cutting activity was observed in 44 randomly selected females from *Prm1*^Tg^; X-C^Tg^ (both X-C-1 and X-C-2 lines), and 14 from *Stra8*^Tg^; X-C-1^Tg^ sires (Suppl. Fig. 4C,D). Sperm from *Prm1*^Tg^; X-C^Tg^ and *Stra8*^Tg^; X-C^Tg^ sires was analysed for indels but none were detected (Suppl. Fig. 4A). One male (out of 20 samples) from a *Ccna1*^Tg^; X-D^Tg^ sire had an indel (Suppl. Fig. 4B), presumably generated by Cas9/gRNA carryover.

**Figure 6.**
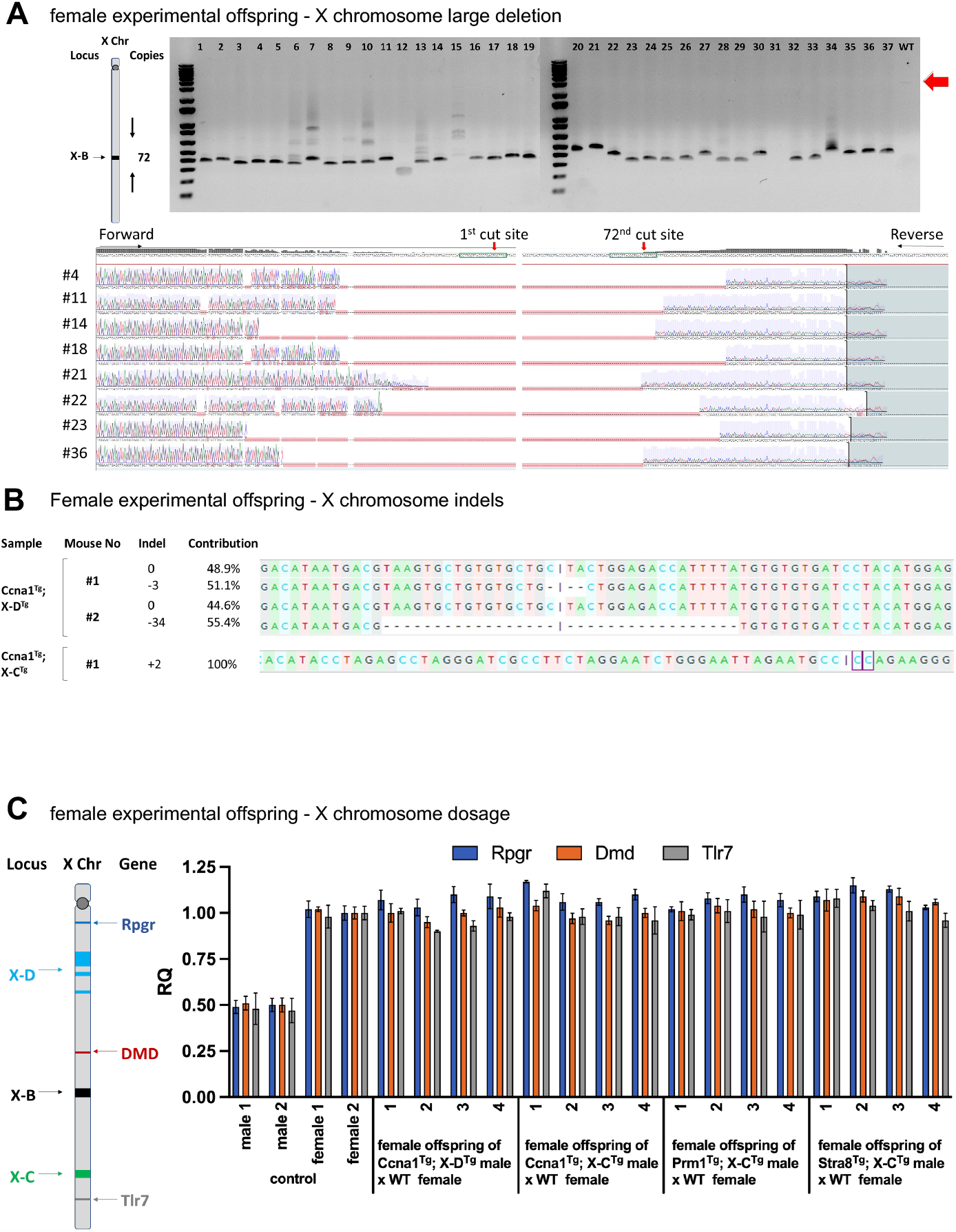
Molecular evidence of X chromosome-targeted Cas9 activity from female offspring of experimental matings. **A**. Schematic of X-B gRNA target region showing number of cuts and spread of cut sites on the mouse X chromosome. PCR across the X-B-targeted cut site region on the X chromosome of pups generated from *Ccna1*^Tg^; X-B^Tg^ male x WT female matings. Expected uncut DNA amplicon size is ∼3000 bp. Smaller products in the DNA gel indicate deletions. Associated Sanger sequencing of X-B gRNA target site from selected samples demonstrates efficient deletion of the region containing repeat sequences. **B**. Sanger sequencing of DNA from female pups of *Ccna1*^Tg^; X-D^Tg^ male x WT female matings (top) and *Ccna1*^Tg^; X-C^Tg^ male x WT female matings (bottom) of X chromosome target sites showing the presence of small indels. **C**. Schematic of X-B, X-C, and X-D target sites and *Rpgr, DMD, Xist*, and *Tlr7* gene loci on the mouse X chromosome. X chromosome dosage qPCR was performed on genomic DNA isolated from female pups of germline promoter-Cas9^Tg^; X shredder^Tg^ male x WT female matings. Expression of *Rpgr, DMD, Xist* and *Tlr7* was normalised to WT female genomic DNA.

Finally, as XO mice are female and viable, we investigated the possibility that female offspring of double transgenic sires did not inherit a paternal X chromosome. X chromosome gene dosage was performed by qPCR analysis of *Rpgr, DMD*, and *Tlr7* in randomly selected female pups from matings of *Ccna1*^Tg^; X-D^Tg^ sire, *Ccna1*^Tg^; X-C^Tg^ sire, *Prm1*^Tg^; X-C^Tg^ sire or *Stra8*^Tg^; X-C^Tg^ sires with WT females. No reduction in X chromosome dosage was detected (Fig. 6C) indicating that paternal X chromosome transmission was not compromised.

## Discussion

Genetic biocontrol technologies for invasive pest and disease vector control hold enormous potential and have progressed significantly in insects. However, it is increasingly clear that translation of molecular strategies from insects to mammals presents significant challenges^3,4^. Here we attempted to develop X-shredder mice, using similar approaches to those employed in *Anopheles* mosquitos, *Drosophila melanogaster* and *Ceratitis capitata*.

As the mechanism that compromises X-bearing gametes in dipteran X-shredders is not completely understood, and a mammalian X-shredder has not previously been developed, the optimal timing for X chromosome cleavage during murine spermatogenesis is not known. We therefore sought to generate Cas9-expressing transgenic lines that were active across different stages of spermatogenesis. The *Stra8* promoter has previously been used to drive transgene expression at premeiotic stages (undifferentiated spermatogonia and preleptotene spermatocytes). Although we successfully generated *Stra8*-Cas9 transgenic mice, expression was very low, approximately 75-fold lower than *Ccna1*-Cas9. No evidence of cleavage or subfertility was detected in *Stra8*-Cas9/gRNA double transgenic lines indicting that Cas9 expression was below the threshold required for experimental analysis. In contrast, the post-meiotic spermatogenic promoter *Prm1* drove high levels of Cas9 mRNA expression in the testes (Fig. 2C). However, no detectable GFP reporter protein was produced (Fig. 2D), and when crossed with X-C gRNA to generate a double transgenic sire, no, or very low frequency, indels were observed in sperm using both X-C-1 and X-C-2 lines (Suppl. Fig. 4A). We note that 3’ UTR regulatory sequences are important for *Prm1* mRNA translation, which may have impacted Cas9 protein levels^17,20^. The *Ccna1* promoter has been shown to drive expression at the initial stages of meiosis (pachytene)^21^. We have previously shown that this *Ccna1*-Cas9 line provides robust Cas9 activity, generating indels in ∼80% of sperm at a single gRNA target site. This transgenic line provides a useful new tool for efficient targeted mutagenesis in male gametes.

Unlike X-shredding dipterans, *Ccna1*-Cas9; gRNA double transgenic male mice did not show significant bias towards male progeny. However, two lines of evidence supported disproportionate loss of X chromosome-bearing sperm, albeit to a small degree; (1) the dosage of X chromosome markers was significantly lower in sperm from some double tg males and (2) *Ccna1*^Tg^; X-D^Tg^ males generated more males than females although this just failed to reach significance.

Instead of male-biased offspring, the most striking impact of germline Cas9 and gRNA expression in double transgenic mice was to block spermatogenesis, manifest as significantly smaller testes volume, lower sperm concentrations, impaired sperm motility, and a reduced number of offspring. Interestingly, triple transgenic mice were even more severely affected with empty tubules (azoospermia) and complete infertility, suggesting that higher levels of X chromosome cleavage resulted in a more severe phenotype. What is the mechanism that underpins this phenotype? Generation of DSBs is a normal feature of meiosis – indeed, hundreds of SPO11-mediated DSBs are generated across autosomes during leptotene which promote homolog recognition and formation of crossovers for meiotic exchange^22,23^. However, the number of DSB is strictly controlled through the calibrated activity of ATM and SPO11 and failure to resolve DSBs inhibits spermatogenesis and results in cell death of pachytene-stage progenitors^24^. Given that (1) DSB formation is normally supressed on the non-homologous regions of the sex chromosomes^25^, (2) leptotene cells are primed for HR repair (rather than NHEJ^26^) and (3) X-chromosome repeat sequences cannot be resolved via synapse formation and crossover due to the absence of a homologous partner, we propose that the spermatogenic block in the *Ccna1*-Cas9; gRNA transgenic males is due, at least in part, to the persistence of unresolved DSBs.

It is interesting to compare the spermatogenic block phenotype in transgenic mice with analogous experiments performed in *Anopheles*. I-PpoI or Cas9 expression in mosquitos is driven by the B-tubulin promoter which is expressed at a similar stage to *Ccna1* in mice. Thus, it appears that spermatogenic checkpoint is less stringent in *Anopheles*, or that the DSBs are resolved more efficiently, or are fewer in number. In any case, it is interesting to note that analysis of mature sperm in I-PpoI transgenic males revealed that the X:Y ratio was similar to WT males^27^. This finding is quite surprising and indicates that male biasing of progeny from this line is due to a post-copulatory effect, likely related to sperm motility, and not via X-shredding. It will be interesting to see what further studies reveal about the mechanism(s) of male biassing in this line, as this information may be also informative for future strategies in mice. And, although we have demonstrated that it is possible to generate competent X-bearing sperm by generating DSB at meiosis (X-B females in Fig. 6A), for the future, it might be more effective to initiate X chromosome cleavage after meiosis to avoid the risk of pachytene spermatogenic block. Indeed, connection of developing spermatids via intercytoplasmic bridges would potentially allow cleavage to occur late in spermatogenesis. Ultimately however, development of a driving-Y version of the X-Shredder would still need to overcome MSCI (which would inhibit transgene expression on the Y), an issue which remains to be solved in insects.

## Methods

### Constructs

The gRNAs (Suppl. Table 1) were cloned into either Cas9 expressing pX459 plasmid, pX458 plasmid or pDG459 plasmid. For the single guide plasmids, a pair of oligonucleotides (Suppl. Table 2) was phosphorylated and ligated into BsbI (NEB) digested pX459 and pX458 plasmids. For dual guide plasmids, two pairs of oligonucleotides (Suppl. Table 2) were phosphorylated and ligated into BsbI (NEB) digested pDG459 plasmid. Corresponding plasmids were sequence verified using single or dual primers (Suppl. Table 2).

### Cell culture and transfection

R1 mouse embryonic stem cells (ES cells) were obtained from Andras Nagy’s laboratory. They were cultured in Dulbecco’s Modified Eagle’s Medium (DMEM; Gibco) supplemented with 20% FCS, 1000 units/ml LIF (ESGRO, Sigma), 3 μM CHIR99021 (Sigma), 1 μM PD0325901 (Sigma), 1 x GlutaMAX, 100 μM non-essential amino acids (Gibco) and 100 μM 2-mercaptoethanol (Sigma). All cells were maintained in humidified incubators at 3711C with 5% CO2 and tested negative for mycoplasma. Plasmids were transfected using Neon transfection system (Invitrogen) following the manufacturer’s protocol. 1 × 10^6^ cells were nucleofected with 10 μg of each plasmid at 1400 V, 10 ms and 3 pulses. Puromycin (2 μg/ml) selection was commenced 24 hrs after nucleofection and continued for two days. Remaining cells were recovered in puromycin free media for another three days followed by cell counting and genomic DNA extraction. All the samples were co-transfected with pX458 plasmid to confirm the transfection efficiency.

### Mouse model generation

Transgenic mice were generated as described in ^28^. The *Ccna1*-Cas9 transgenic line used was described previously^6^.

### Blastocyst generation

Fertilised embryos were generated using IVF as described at https://card.medic.kumamoto-u.ac.jp/card/english/sigen/manual/lowtempce.html. Fertilised embryos (observed to have two pronuclei) were cultured under oil to blastocyst and collected in 1μl of handling media into 9μl of blastocyst lysis buffer and frozen.

### DNA extraction

Genomic DNA was extracted from mouse tail tip or ear notch biopsies using the High Pure PCR Template Preparation Kit (Roche) following the manufacturer’s protocol. A pair of primers (Suppl. Table 2) were used to amplify the targeted fragment which covering a single cut site or multiple cut sites on ChrX. PCR products were purified by using QIAquick PCR Purification Kit, Qiagen. Sanger sequenced samples were analysed using both ICE and Decodr tools (https://ice.synthego.com/#/ and https://decodr.org/, respectively) to detect the indels at target sites.

Blastocysts were lysed in 10μl (final volume – see above) blastocyst lysis buffer (tRNA from baker’s yeast, Proteinase K, and blastocyst lysis buffer base; Tris-HCl, KCl, Gelatin, Tween 20, H_2_0), placed in a thermocycler for 1 cycle of 56°C for 10 min, and 95°C for 10 min followed by 4°C. PCR was performed using 4 μl of the mixture and a pair of oligos (Suppl. Table 2) to amplify the target fragment which covered a single cut site or multiple cut sites on the X chromosome.

Sperm gDNA extraction was performed as previously described^5^.

### Genotyping

Genomic DNA was extracted from ear punch biopsies using the Monarch Genomic DNA purification kit (NEB). Transgenes were identified using PCR primers specific for each transgene (Suppl. Table 2) amplified using Taq DNA polymerase (Roche) and Failsafe Buffer D (Epicentre). PCR products were run on a 1% agarose gel and imaged using a Gel Doc XR+ (Bio-Rad).

### Digital PCR

150 ng of genomic DNA extracted from ear punch biopsies (see Genotyping protocol) was digested with MseI (NEB) and 1x CutSmart buffer at 37°C for two hours. The ThermoFisher Scientific QuantStudio 3D protocol for digital PCR was followed. In brief, digested genomic DNA was diluted and PCR performed with QuantStudio 3D Digital PCR master mix v2 and probes against Rpp30, GFP, and mCherry. The following thermal cycle was used: Initial denaturation 96°C for 10 mins, annealing 60°C for 2 mins, extension 98°C for 30 secs, 39 cycles, final extension 60°C for 2 mins, hold at 10°C. Digital PCR chips were read and analysed with ThermoFisher Scientific AnalysisSuite software.

### RNA extraction

Testis were harvested from mice and stored in 1.5 mL tubes at -80°C until the day of processing. Testis were thawed on ice and 500 μL of TRIzol was added, then the testis was transferred to a small dish and cut into 2 mm pieces using a scalpel blade. The tissue was transferred back to the 1.5 mL tube and placed on ice while additional samples were processed. Homogenised tissue samples were incubated for 5 mins at RT, before 100 μL of chloroform was added. Tubes were vortexed followed by incubation for 2 mins at RT and centrifugation for 15 mins at 12,000 rcf at 4°C. The aqueous upper layer (∼175 μL) was transferred to a new 1.5 mL tube, 100 μL of 70% ethanal added and the sample mixed by pipetting. The RNAeasy extraction kit (Qiagen) was used for RNA purification. In brief, 700 μL of sample was added to an RNeasy Mini Spin Column in a collection tube (Qiagen) and centrifuged for 30 secs at 8,000 rcf. The flow through was discarded and the column placed back in the collection tube. 350 μL of Buffer RW1 was added and centrifuged for 30 secs at 8,000 rcf. A mix of 70 μL of Buffer RDD (Qiagen) and 10 μL DNase I stock solution (Qiagen) was then added to the column and incubated for 15 mins at RT. 350 μL Buffer RW1 was added followed by centrifugation for 30 secs at 8,000 rcf. The flow through was discarded and the column placed back in the collection tube. 500 μL Buffer RPE was added followed by centrifugation for 2 mins at 8,000 rcf. The flow through was discarded and the column was added to a new collection tube. The column was spun for 1 min at 18,000 rcf, then transferred to a labelled 1.5 mL tube. 30 μL of RNase-free H_2_O was added to the column membrane, incubated for 1 min at RT, then centrifuged for 1 min at 8,000 rcf.

### qRT-PCR

cDNA was generated using the High-Capacity RNA-to-cDNA kit (Applied Biosystems) whereby 2 ug of RNA was added to 10 μL of 2x RT Buffer Mix and 1 uL of 20x RT Enzyme Mix and made up to a total volume of 20 μL using nuclease-free H_2_O. The tubes were placed in a thermocycler with the following parameters: 37°C for 60 mins, 95°C for 5 mins, 4°C hold. qPCR was performed with Fast SYBE Green Master Mix, qPCR primers as described in Supp Table 2, and 0.5 μL of cDNA. The qPCR reaction was run on a QuantStudio 3 (Applied Biosystems) and data analysed with Design and Analysis 2.7.0 software (Applied Biosystems).

### Collection of sperm

Sperm were collected post mortem from the cauda epididymidis and ductus deferens and expressed into 1ml of G-IVF medium (Vitrolife, Gothenberg, Sweden) and incubated for at least 10 min in 6% CO2 at 37°C. Sperm concentration and motility were measured on the CASA® semi-automatic semen analyser (Microptic, Spain, Barcelona), where at least 500 sperm were counted across a minimum of 5 fields of view. Low-and high-quality control beads (Microptic) were run prior to each sample analysis. A pre-set mouse count/motility program was used to calculate sperm concentration, rapid progressive motility (>25 μm/s), medium progressive motility (<15 μm/s), non-progressive (<10 μm/s) and immotile sperm. Remaining isolated sperm (∼990 μl) were washed in sterile PBS at 400g for 5 min x 2, snap frozen and stored at -80oC until further analysis.

### qPCR

To assess the X chromosome dosage, gDNA was extracted from biopsies and 50 ng used in the assay. Gene-specific primers for Rpgr, DMD, and Tlr7 (Suppl. Table 2) were used together with fast SYBR Green quantitative real-time PCR (qPCR) reagent. The mouse Sox1 gene was amplified using gene specific primers as an internal control gene. The thermal cycling conditions included one cycle at 95°C for 20 s, followed by 40 cycles of 95°C for 1 s, and 60°C for 20 s. Final single cycles at 95°C for 1 s, 60°C for 20 s, and 95°C for 1 s were carried out to assess the specificity of the target amplification. The Livak method^29^ was used to quantify the level of transcripts.

### Histology

Mice were euthanized by CO_2_ inhalation, testes were dissected and fixed in 4% paraformaldehyde overnight at 4°C. Tissues were washed with PBS and incubated overnight in 30% sucrose at 4°C. OCT embedded tissues were stored at -80°C. Embedded tissues were cut into 8 μm sections and routine hematoxylin and eosin (H&E) staining was performed. Slides were quantified for the percentage of tubules with normal spermatogenesis as outlined by^30^, with at least 200 seminiferous tubules across four sections assessed for each mouse.

### Immunofluorescence

Transfected cells were cultured on the gelatin coated coverslips for 24 hrs. Cells were fixed in 4% paraformaldehyde for 15 min and stained with anti-γ-H2AX (CST,9718) and chicken anti-GFP (Abcam, #ab13970) primary antibodies in 1:400 and 1:600 dilution, respectively. An anti-rabbit 594 (Thermo Fisher Scientific, #A-11012) and anti-chicken 488 (Thermo Fisher Scientific, #A-11039) secondary antibodies were used in 1:300 dilution. The fluorescence images were taken on Nikon microscope using the same settings across the samples. H2AX and GFP double positive cells were randomly counted.

### Animal ethics

All animal work was conducted in accordance with Australian guidelines for the care and use of laboratory animals following approval by the SAHMRI Animal Ethics Committee (approval number SAM439.19).

## Figure Legends

**Supplementary Figure 1.** Molecular evidence of X chromosome-targeted Cas9 activity in mouse ES cells. Genomic DNA from the surviving cells was isolated and a single X-C gRNA target site and X-D gRNA target site was amplified and sequenced. The presence of indels is shown in either single gRNA or dual gRNA plasmid transfected cells.

**Supplementary Figure 2.** Transgene construct design. **A**. Murine germline-specific promoters *Ccna1, Prm1*, and *Stra8* were inserted upstream of an NLS-flanked Cas9 coding sequence followed by the self-cleaving linker, P2A, EGFP, WPRE and a bGH poly(A). **B**. Expression of X shredder gRNAs was driven by the human U6 promoter and supported by the U6 3’ regulatory sequence downstream of the guide scaffold sequence. In the opposite direction, mCherry expression was driven by CMV enhancer and CMV promoter sequence. **C**. Schematic of sperm development stage with *Ccna1*-Cas9-EGFP construct expression indicated as weak (light green) and strong (dark green) boxes.

**Supplementary Figure 3.** Fertility impacts of X shredder activity. Litter size (left) and pregnancy success as determined by plug and successful pregnancy counts (right) were performed. **A**. *Ccna1*^Tg^; X-B^Tg^ x WT matings. Single Tg n = 8, *Ccna1*^Tg^; X-B^Tg^ dam n = 10, *Ccna1*^Tg^; X-B^Tg^ sire n = 22. **B**. *Ccna1*^Tg^; X-C^Tg^ x WT matings (two independent X-C^Tg^ founder lines). Single Tg n = 9, *Ccna1*^Tg^; X-C-1^Tg^ dam n = 3, *Ccna1*^Tg^; X-C-1^Tg^ sire n = 15, *Ccna1*^Tg^; X-C-2^Tg^ dam n = 3, *Ccna1*^Tg^; X-C-2^Tg^ sire n = 0. **C**. *Prm1*^Tg^; X-C^Tg^ x WT matings (two independent X-C^Tg^ founder lines). Single Tg n = 6, *Prm1*^Tg^; X-C-1^Tg^ sire n = 14, *Prm1*^Tg^; X-C-2^Tg^ sire n = 27. **D**. *Stra8*^Tg^; X-C^Tg^ x WT matings (two independent X-C^Tg^ founder lines). Single Tg n = 4, *Stra8*^Tg^; X-C-1^Tg^ sire n = 10, *Stra8*^Tg^; X-C-2^Tg^ sire n = 18. **E**. *Ccna1*^Tg^; X-D^Tg^ x WT matings. Single Tg n = 12, *Ccna1*^Tg^; X-D^Tg^ dam n = 8, *Ccna1*^Tg^; X-D^Tg^ sire n = 50.

**Supplementary Figure 4.** Sex of blastocysts from *in vitro* fertilisation using sperm from *Ccna1*^Tg^; X-D^Tg^ males. **A**. PCR for the Y-chromosomal *Sry* gene in DNA extracted from blastocysts generated using *Ccna1*^Tg^; X-D^Tg^ sperm and WT oocytes. **B**. Sanger sequencing of a single X-D gRNA target site on the X chromosome.

**Supplementary Figure 5.** Molecular characterisation of X shredder transgenic male sperm and X chromosome target sites in female offspring from experimental matings. **A**. Sperm from ex-breeder X-C^Tg^, *Prm1*^Tg^; X-C^Tg^ and *Stra8*^Tg^; X-C^Tg^ males was isolated and Sanger sequencing performed for the respective X chromosome target sites to identify the presence of indels. **B**. Genomic DNA of a male pup from a *Ccna1*^Tg^; X-D^Tg^ male x WT female mating was isolated and a single X-D gRNA target site was sequenced. The presence of a small proportion of indels is shown. **C**. Genomic DNA of female pups from a *Prm1*^Tg^; X-C^Tg^ male x WT female, *Stra8*^Tg^; X-C^Tg^ male x WT female, *Ccna1*^Tg^; X-C^Tg^ male x WT female, and *Ccna1*^Tg^; X-D^Tg^ male x WT female mating was isolated and a single X-C gRNA target site was sequenced. No indels were detected in any of the analysed female offspring.

