## Supplementary Figures for "Investigating the potential of X shredding for mouse genetic biocontrol"

### Slide 1
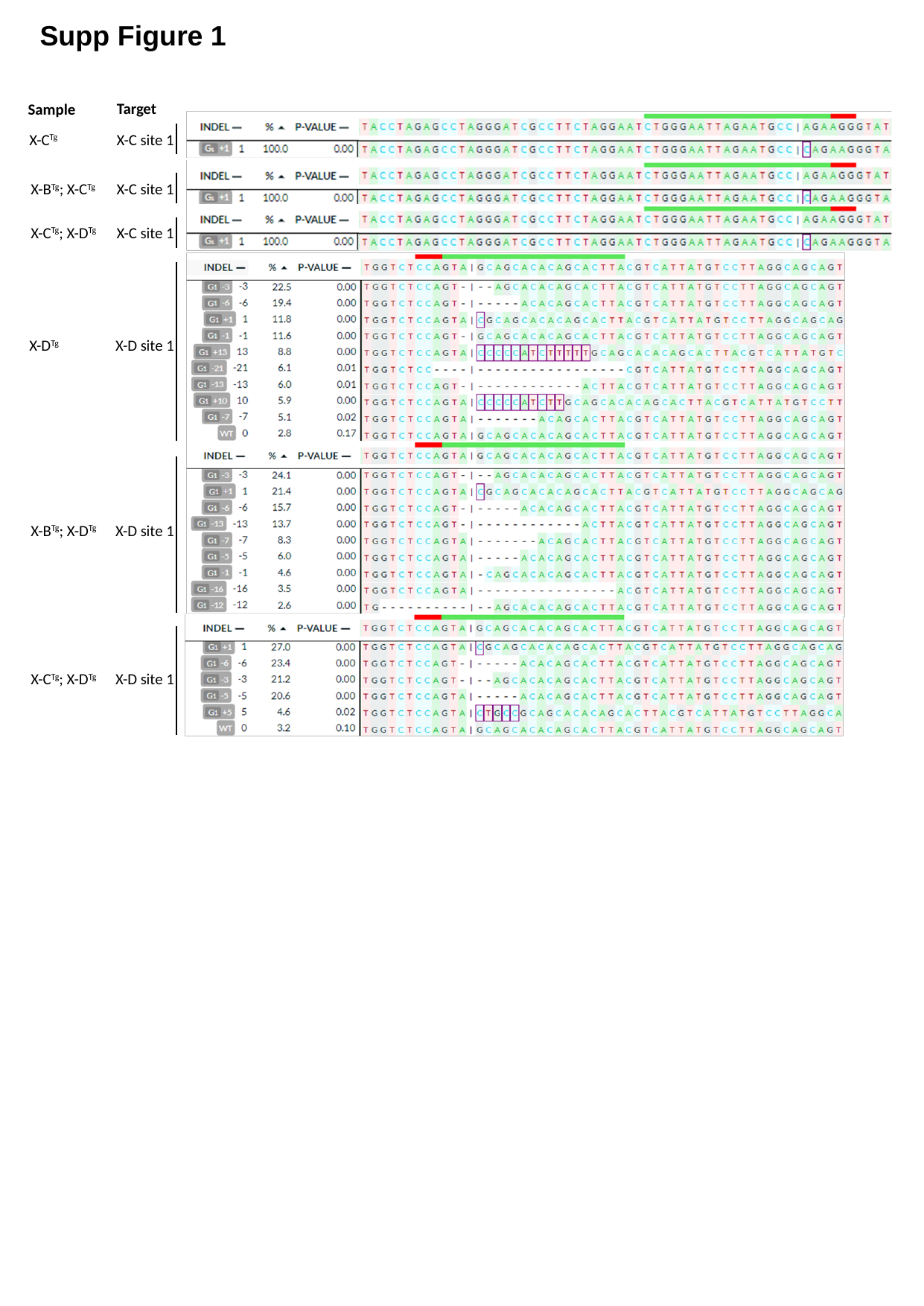

Supp Figure 1
Target
Sample
X-C^Tg^
X-C site 1
X-B^Tg^; X-C^Tg^
X-C site 1
X-C^Tg^; X-D^Tg^
X-C site 1
X-D^Tg^
X-D site 1
X-B^Tg^; X-D^Tg^
X-D site 1
X-C^Tg^; X-D^Tg^
X-D site 1

### Slide 2
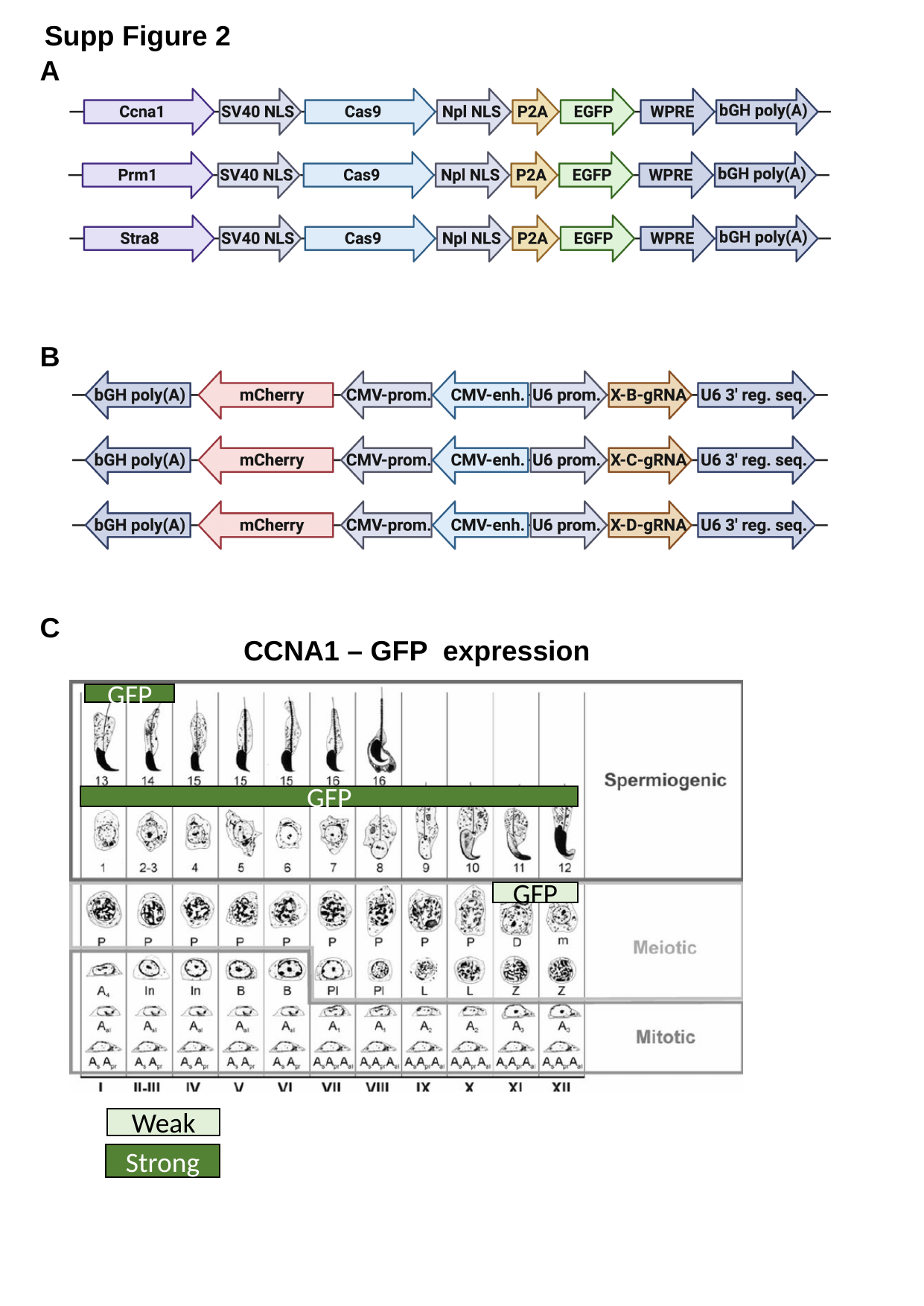

Supp Figure 2
A
B
C
CCNA1 – GFP expression
GFP
GFP
GFP
Weak
Strong

### Slide 3
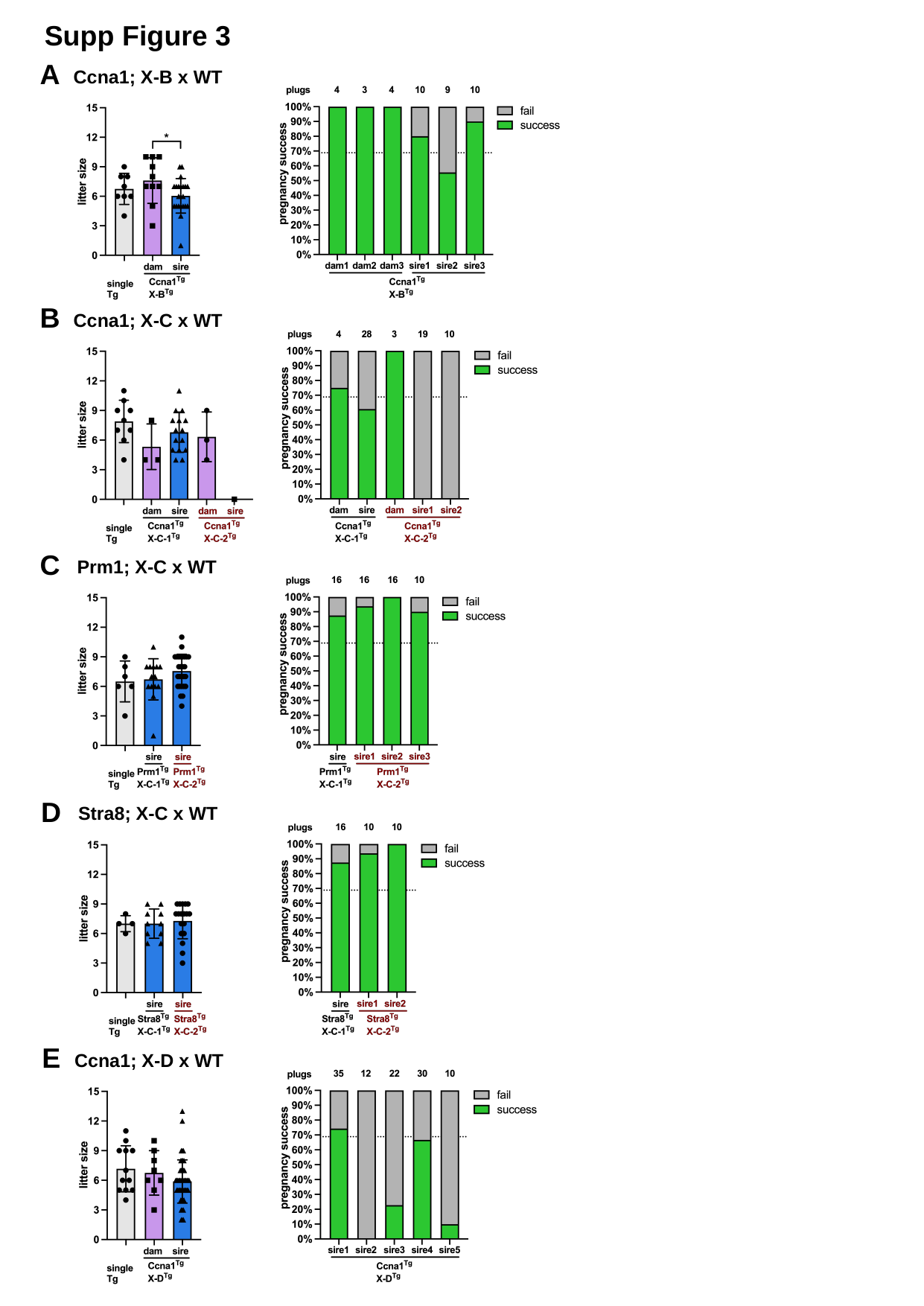

Supp Figure 3
A
Ccna1; X-B x WT
B
Ccna1; X-C x WT
C
Prm1; X-C x WT
D
Stra8; X-C x WT
E
Ccna1; X-D x WT

### Slide 4
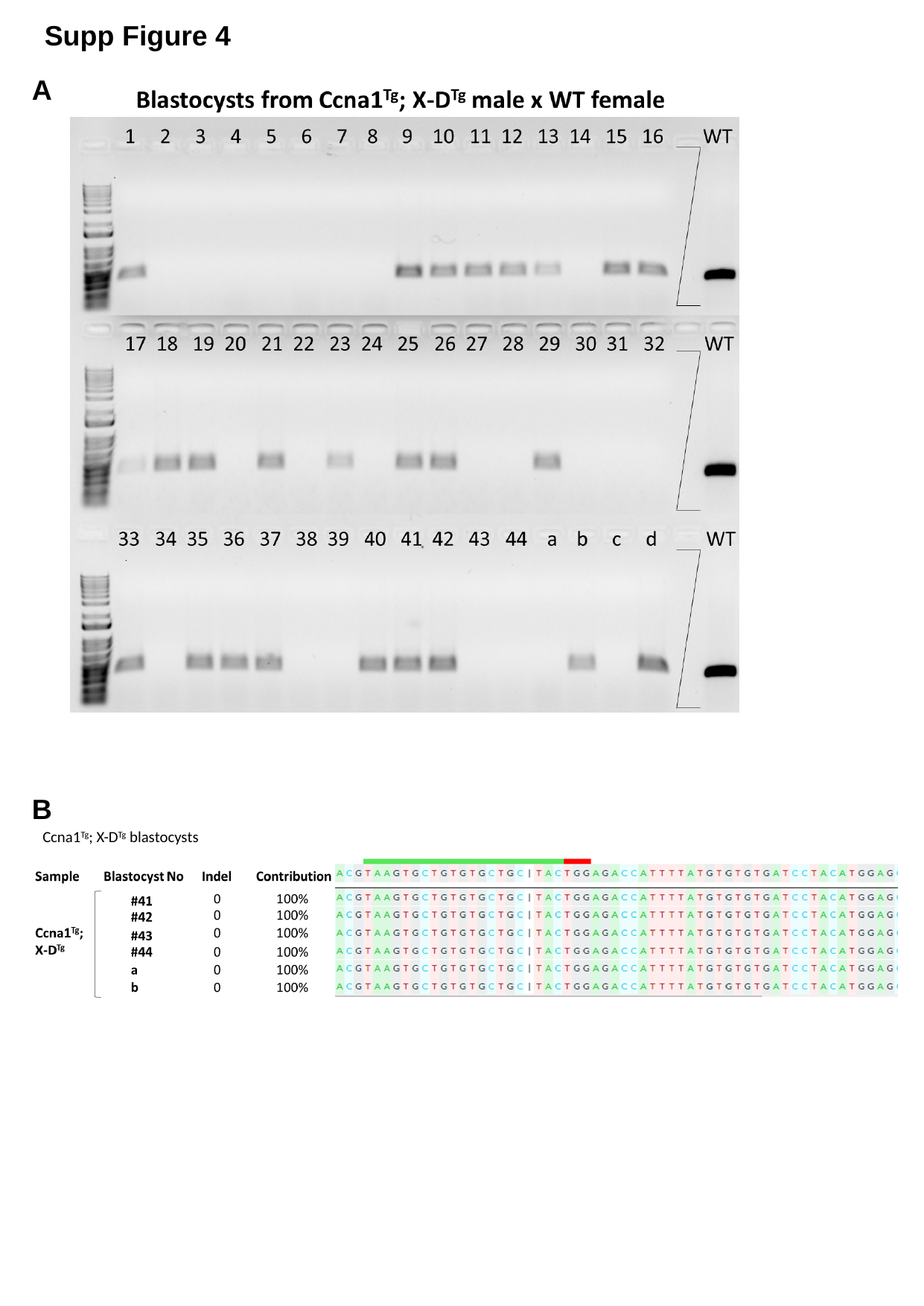

Supp Figure 4
A
B
Ccna1^Tg^; X-D^Tg^ blastocysts

### Slide 5
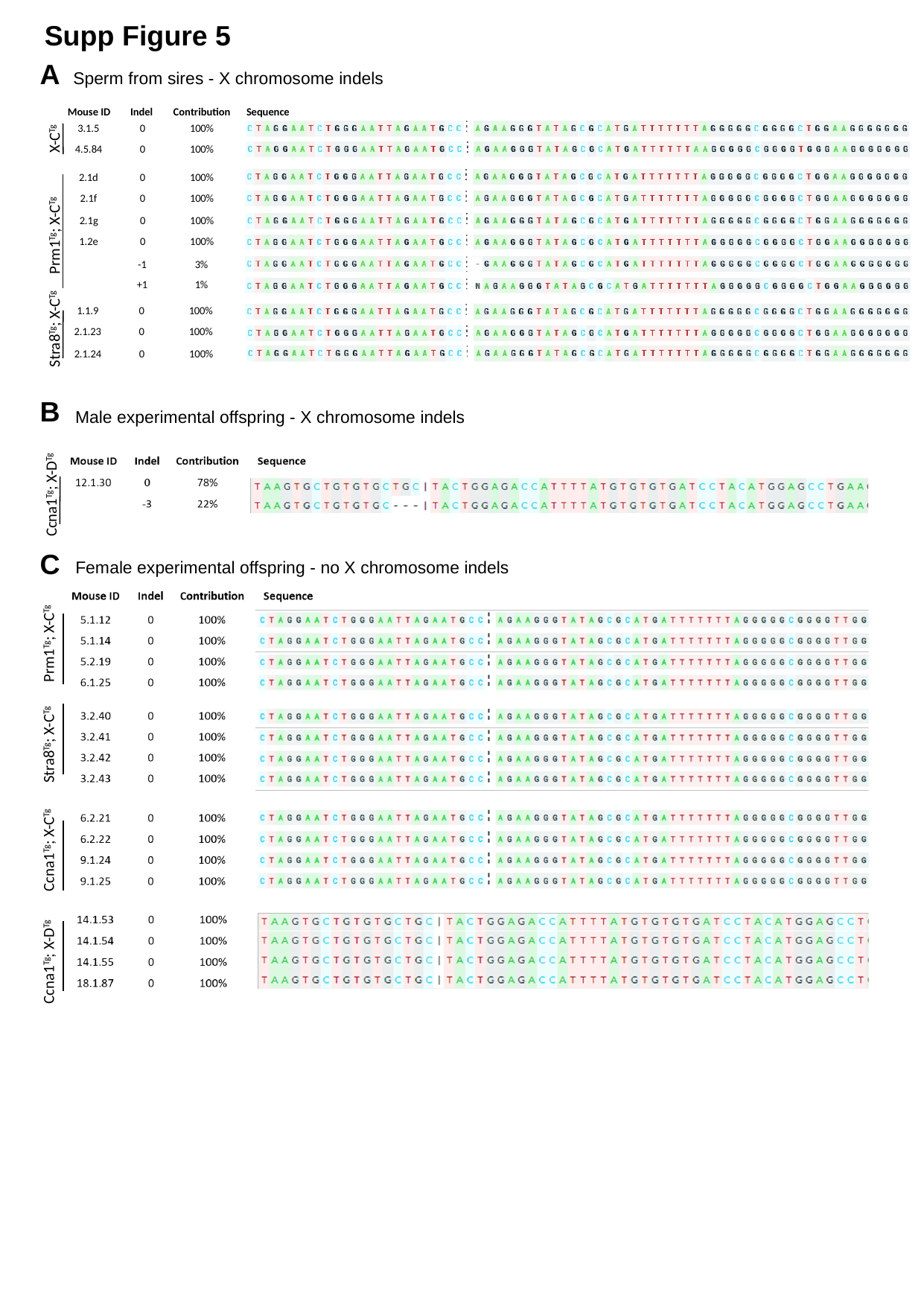

Supp Figure 5
A
Sperm from sires - X chromosome indels
Sequence
Mouse ID
Indel
Contribution
3.1.5
0
100%
4.5.84
0
100%
2.1d
0
100%
2.1f
0
100%
2.1g
0
100%
1.2e
0
100%
-1
3%
+1
1%
1.1.9
0
100%
2.1.23
0
100%
2.1.24
0
100%
X-C^Tg^
Prm1^Tg^; X-C^Tg^
Stra8^Tg^; X-C^Tg^
B
Male experimental offspring - X chromosome indels
Ccna1^Tg^; X-D^Tg^
C
Female experimental offspring - no X chromosome indels
Prm1^Tg^; X-C^Tg^
Stra8^Tg^; X-C^Tg^
Ccna1^Tg^; X-C^Tg^
Ccna1^Tg^; X-D^Tg^

### Slide 6
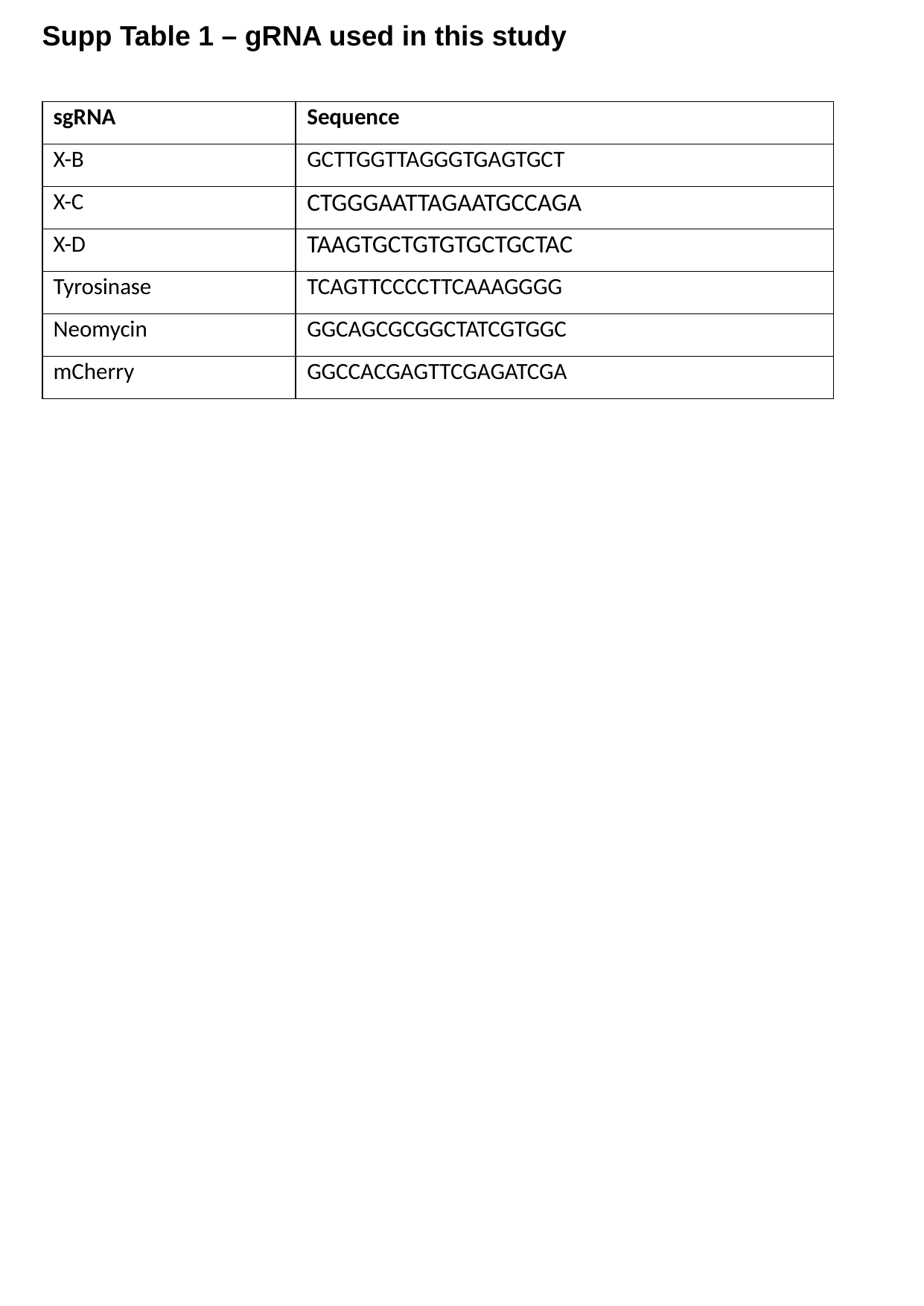

Supp Table 1 – gRNA used in this study
| sgRNA | Sequence |
| --- | --- |
| X-B | GCTTGGTTAGGGTGAGTGCT |
| X-C | CTGGGAATTAGAATGCCAGA |
| X-D | TAAGTGCTGTGTGCTGCTAC |
| Tyrosinase | TCAGTTCCCCTTCAAAGGGG |
| Neomycin | GGCAGCGCGGCTATCGTGGC |
| mCherry | GGCCACGAGTTCGAGATCGA |

### Slide 7
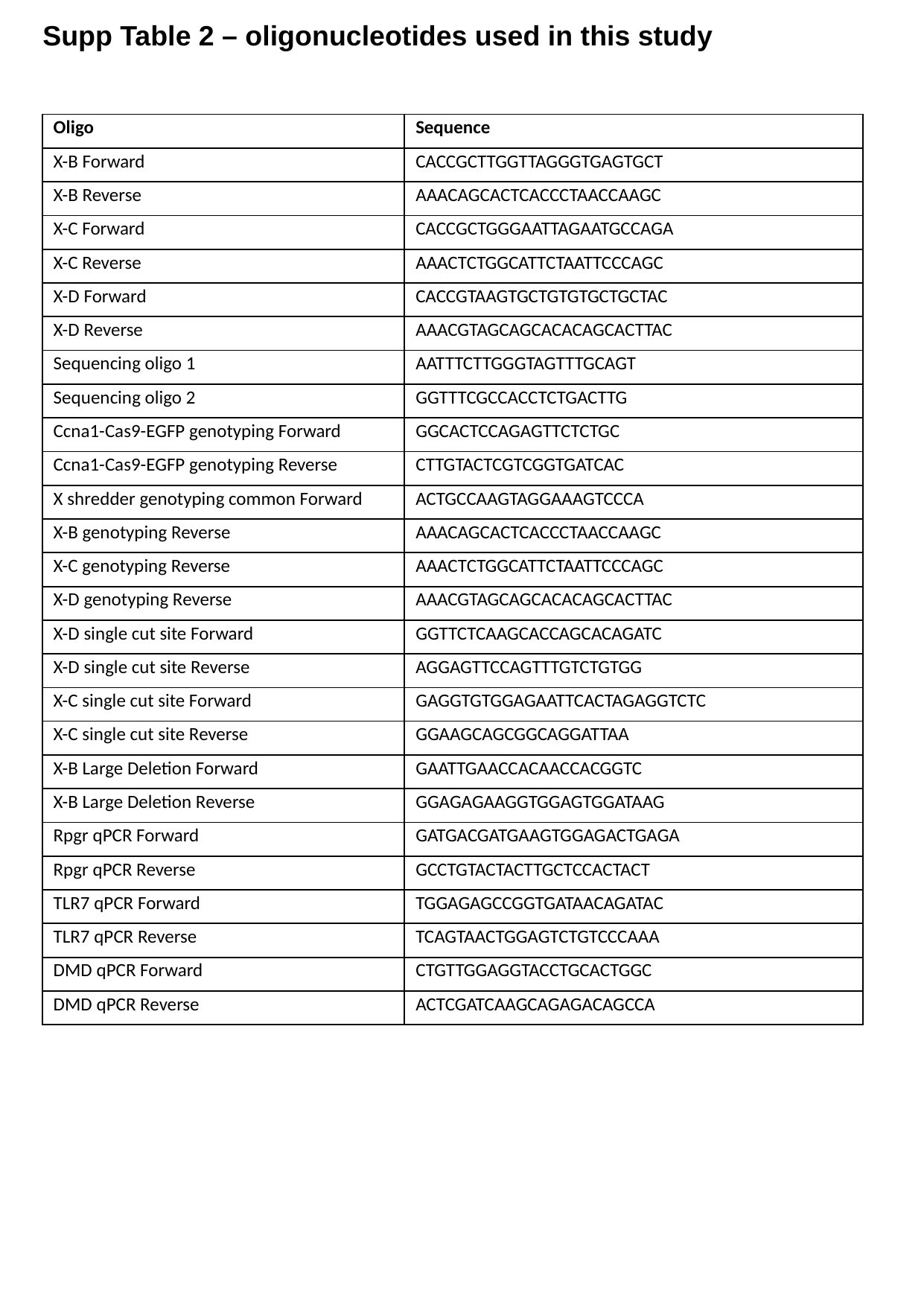

Supp Table 2 – oligonucleotides used in this study
| Oligo | Sequence |
| --- | --- |
| X-B Forward | CACCGCTTGGTTAGGGTGAGTGCT |
| X-B Reverse | AAACAGCACTCACCCTAACCAAGC |
| X-C Forward | CACCGCTGGGAATTAGAATGCCAGA |
| X-C Reverse | AAACTCTGGCATTCTAATTCCCAGC |
| X-D Forward | CACCGTAAGTGCTGTGTGCTGCTAC |
| X-D Reverse | AAACGTAGCAGCACACAGCACTTAC |
| Sequencing oligo 1 | AATTTCTTGGGTAGTTTGCAGT |
| Sequencing oligo 2 | GGTTTCGCCACCTCTGACTTG |
| Ccna1-Cas9-EGFP genotyping Forward | GGCACTCCAGAGTTCTCTGC |
| Ccna1-Cas9-EGFP genotyping Reverse | CTTGTACTCGTCGGTGATCAC |
| X shredder genotyping common Forward | ACTGCCAAGTAGGAAAGTCCCA |
| X-B genotyping Reverse | AAACAGCACTCACCCTAACCAAGC |
| X-C genotyping Reverse | AAACTCTGGCATTCTAATTCCCAGC |
| X-D genotyping Reverse | AAACGTAGCAGCACACAGCACTTAC |
| X-D single cut site Forward | GGTTCTCAAGCACCAGCACAGATC |
| X-D single cut site Reverse | AGGAGTTCCAGTTTGTCTGTGG |
| X-C single cut site Forward | GAGGTGTGGAGAATTCACTAGAGGTCTC |
| X-C single cut site Reverse | GGAAGCAGCGGCAGGATTAA |
| X-B Large Deletion Forward | GAATTGAACCACAACCACGGTC |
| X-B Large Deletion Reverse | GGAGAGAAGGTGGAGTGGATAAG |
| Rpgr qPCR Forward | GATGACGATGAAGTGGAGACTGAGA |
| Rpgr qPCR Reverse | GCCTGTACTACTTGCTCCACTACT |
| TLR7 qPCR Forward | TGGAGAGCCGGTGATAACAGATAC |
| TLR7 qPCR Reverse | TCAGTAACTGGAGTCTGTCCCAAA |
| DMD qPCR Forward | CTGTTGGAGGTACCTGCACTGGC |
| DMD qPCR Reverse | ACTCGATCAAGCAGAGACAGCCA |
